# Methylation QTLs are associated with coordinated changes in transcription factor binding, histone modifications, and gene expression levels

**DOI:** 10.1101/006171

**Authors:** Nicholas E. Banovich, Xun Lan, Graham McVicker, Bryce van de Geijn, Jacob F. Degner, John D. Blischak, Julien Roux, Jonathan K. Pritchard, Yoav Gilad

## Abstract

DNA methylation is an important epigenetic regulator of gene expression. Recent studies have revealed widespread associations between genetic variation and methylation levels. However, the mechanistic links between genetic variation and methylation remain unclear. To begin addressing this gap, we collected methylation data at ∼300,000 loci in lymphoblastoid cell lines (LCLs) from 64 HapMap Yoruba individuals, and genome-wide bisulfite sequence data in ten of these individuals. We identified (at an FDR of 10%) 13,915 *cis* methylation QTLs (meQTLs)—i.e., CpG sites in which changes in DNA methylation are associated with genetic variation at proximal loci. We found that meQTLs are frequently associated with changes in methylation at multiple CpGs across regions of up to 3 kb. Interestingly, meQTLs are also frequently associated with variation in other properties of gene regulation, including histone modifications, DNase I accessibility, chromatin accessibility, and expression levels of nearby genes. These observations suggest that genetic variants may lead to coordinated molecular changes in all of these regulatory phenotypes. One plausible driver of coordinated changes in different regulatory mechanisms is variation in transcription factor (TF) binding. Indeed, we found that SNPs that change predicted TF binding affinities are significantly enriched for associations with DNA methylation at nearby CpGs.

**Author Summary:** DNA methylation is an important epigenetic mark that contributes to many biological processes including the regulation of gene expression. Genetic variation has been associated with quantitative changes in DNA methylation (meQTLs). We identified thousands of meQTLs using an assay that allowed us to measure methylation levels at around 300 thousand cytosines. We found that meQTLs are enriched with loci that is also associated with quantitative changes in gene expression, DNase I hypersensitivity, PolII occupancy, and a number of histone marks. This suggests that many molecular events are likely regulated in concert. Finally, we found that changes in transcription factor binding as well as transcription factor abundance are associated with changes in DNA methylation near transcription factor binding sites. This work contributes to our understanding of the regulation of DNA methylation in the larger context of gene regulatory landscape.

## Introduction

Changes in gene expression levels are important contributors to phenotypic variation in human populations [1–11]. One way in which gene expression levels may be altered is through changes in chromatin function [12–19]. Recent studies have focused on identifying genetic variants that impact chromatin function [14,20] by studying inter-individual variation in DNase I sensitivity, a general indicator of chromatin accessibility [21], as well as a variety of histone modifications [17–19,22]. A single genetic variant was often found to be associated with coordinated changes in multiple molecular phenotypes, including chromatin accessibility, nucleosome positioning, chromatin modifications and gene expression levels [17–19]. In many cases of coordinated changes, the associated genetic variants seem to act through the disruption of transcription factor binding sites [17–19]. This body of work highlights the value of using multiple molecular phenotypes to understand the connection between genetic variation and gene expression. One important epigenetic mark not considered by these recent integrated studies is DNA methylation.

DNA methylation refers to the addition of a methyl group to cytosine nucleotides. In vertebrates, DNA methylation primarily affects cytosines that are immediately 5’ to guanines, i.e., CpGs. Appropriate methylation is essential for development and cellular differentiation [23–25]. Changes in DNA methylation levels have been linked to a number of diseases including tumorigenesis, [26,27] age-related defects [28,29] and mental disorders [30,31]. Typical array-based methylation assays provide a single measurement for each CpG site, which is interpreted to reflect the proportion of cells in which a given site is methylated. In general, this measurement was found to have a bimodal distribution across sites [12,32-34], which is believed to indicate that most sites are either methylated or unmethylated in nearly all cells in a given tissue or culture. Some measurements, however, are intermediate [33] (we refer to these as ‘intermediate methylation levels’), which could either reflect methylation in a subset of cells or just in a single allele (one chromosome) in each cell. Most unmethylated CpGs are within CpG islands (CGIs), namely regions in the genome in which many CpGs are located in close proximity [33,35,36]. CGIs account for a small proportion of CpGs in the genome but they tend to be located near transcription start sites (TSSs). The methylation levels of CGIs are generally negatively correlated with the expression levels of nearby genes [12,33,35-37], an observation that led to a common early belief that DNA methylation was primarily a repressive epigenetic mark.

A number of studies have shown that genetic variation is often associated with quantitative changes in methylation levels [8,12,38-40]. Early QTL studies focused on methylation data from relatively few CpGs with a heavy bias towards promoter regions. A more recent study that used a comprehensive array platform considered genome-wide patterns and reported over 20,000 methylation QTLs (meQTLs [40]). A number of meQTLs were also shown to be associated with changes in gene expression level (namely, these meQTLs are also classified as eQTLs) [8,12,40], although it is not clear whether the methylation changes are a cause or consequence of the gene expression changes [40]. Interestingly, in contrast to the early belief that methylation is primarily associated with repression, both direct and inverse correlations between methylation and gene expression levels have been observed. This suggests that the relationship between DNA methylation and gene expression levels may depend on the genomic context of the CpG [8,12,40].

In general, the mechanisms by which DNA methylation levels are being regulated remain unclear. One likely pathway is through coordination between DNA methylation and chromatin modifiers. For example, H3K4 methyltransferase is recruited by CFP1, which binds to unmethylated CpG islands [41]. In turn, H3K27me3 and DNA methylation have been shown to have mutually exclusive gene silencing functions, in at least some cases [42,43]. There is also limited evidence that TF binding may be associated with nearby changes in DNA methylation. For example, the insertion of a CTCF binding site was shown to cause changes in methylation levels near the insertion site (presumably due to the binding of CTCF) [34,44]. Less direct evidence comes from observations that TF binding sites are enriched in differentially methylated regions (DMRs) between individuals and cell types [45]. However, it is still unclear how frequently changes in TF binding affect the DNA methylation levels of nearby CpGs. It is also unclear whether this is a property that is associated with the binding of most TFs or only a selected few. More generally, there has not yet been a broad examination of coordination between meQTLs and other molecular phenotypes.

In the current study, we therefore examined associations and correlations between genetic variation, DNA methylation, and multiple additional cellular regulatory phenotypes. We focused on a panel of Yoruba HapMap lymphoblastoid cell lines (LCLs), which have been extensively characterized in previous work. In addition to the methylation data we collected for the present study, genomic sequences are available for the majority of these lines [21], as well as RNA sequencing data and DNase I sensitivity profiles [21]. Histone modification data (profiles for H3K4mel, H3K4me3, H3K27ac, H3K27me3P) and PolII ChIP-seq data are also available for a subset of these lines [19].

## Results

We measured methylation levels in 64 Yoruba LCLs using the Illumina Infinium HumanMethylation450 array, which assays methylation levels at roughly 450,000 cytosines, the majority of which are in CpGs. Probes on this array particularly target CpGs near transcription start sites, including CpG islands and CpG shores. As a first step in our data processing, we excluded array probes that did not uniquely map to the human genome as well as probes that overlapped a known sequence variant (see Methods). After these filtering steps we retained methylation measurements from 340,485 probes. As was suggested in previous studies [40,46,47], we quantile-normalized the data to a standard normal within each individual and across probes (though we considered the effects of alternative normalization approaches; see Methods). To account for unobserved confounders we performed principal component analysis. We found that removing four principal components maximized our power to identify meQTLs. Further details on the data processing, normalization, and tests for the effect of confounders are provided in the Methods. In addition to the array data from 64 individuals, we also collected low-coverage whole-genome bisulfite sequencing data from a subset of ten individuals (median genomic coverage 2.4x; see Methods).

## Mapping methylation QTLs

We first examined the association between genetic variation and differences in methylation levels across individuals. For this analysis, we considered only the array data (because we performed whole-genome bisulfite sequencing in only ten individuals). We used previously collected and imputed [21] genotype data for the 64 individuals from the HapMap and 1000 Genomes Projects [48,49]. We focused on proximal (putatively *cis*) associations between genotypes and DNA methylation levels by considering, in each case, genetic variation within a 6 kb region centered on the genomic location of a methylation probe on the array. This window size was chosen because smaller and larger windows yielded fewer significant associations at a given FDR. At an FDR of 10% we identified 13,915 CpG sites with at least one *cis* meQTL (Fig. 1A). When multiple SNPs were significantly associated with methylation levels at a given site, we only considered (for the purpose of counting the overall number of meQTLs) the single most significant association. Since the methylation data measured by nearby pairs of probes are frequently correlated, we wondered whether this analysis might overstate the number of independent meQTL signals. To address this, we examined pairwise correlations of data from all probes located within 5 kb of each other. We found that data from only 203 or 520 of the associated probes (normalized or untransformed data, respectively) are significantly correlated (Pearson Correlation and a T-test; *P* < 0.05) suggesting that the reported number of independent meQTL is not substantially inflated by correlation of the methylation data across nearby probes.

**Figure 1.**
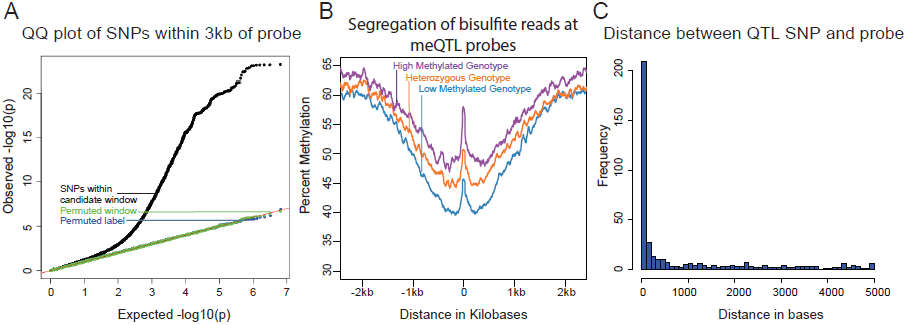
**A)** QQ plot of –log10 p-values for testing the null of no association between methylation levels measured by all probes that passed our quality filters, and all SNPs within 3 kb of these probes. Data for SNPs within the candidate window are in black; negative control SNPs for which we chose a random 6 kb window elsewhere in the genome are in green; SNPs with the genotype labels permuted are in blue. **B)** Average methylation levels estimated using the bisulfite sequence data at meQTL probes, segregated by meQTL genotype. **C)** Histogram showing the distribution of distances between meQTL SNPs and the associated methylated sites in base pairs, for meQTLs where there is a single most likely causal site.

We next used the genome-wide bisulfite sequencing data to provide a general validation of meQTL associations that were identified using the array data (Fig. 1B), as well as to investigate whether meQTLs are generally associated with changes in methylation at a single CpG or a larger region. In general, we observed a high correlation between the estimates of methylation levels based on the array data and the estimates of methylation levels based on the whole genome bisulfite sequencing (R = 0.93; Fig. S1). We note that the read depth and sample size of the bisulfite sequencing data set are insufficient to allow for validation of individual meQTL. Instead, we aggregated the sequence data by considering the centers of probe locations whose methylation data are associated with meQTLs (see Methods for more details). Using that approach, we found a clear difference in methylation level across meQTL genotypes. In addition, we observed a broad-scale association of meQTL genotypes with methylation levels over a region extending between 1.5 and 2 kb in either direction from the methylation loci originally probed by the array. This result indicates that multiple CpGs within a local region are often associated with a single meQTL.

We sought to estimate the typical distance between meQTLs and the location of associated methylated sites (based on the genomic location of the array probes). This analysis is complicated by the fact that, due to LD, it is often unclear which site is causal for any given meQTL. We thus focused on a subset of associations that are more likely to be causal, namely on 409 meQTLs that are the only strongly associated loci within 5 kb of the methylated site (see Methods). Our approach does not provide direct evidence that these are indeed causal sites, but without additional experimental data (namely, using only the meQTL mapping framework), it is likely the best approach to obtain a subset of loci that is enriched with true causal associations [7,11]. These 409 meQTLs are generally located very near the associated methylation site (the median distance is 76bp; Fig. 1C), with only 52 (13%) of the putatively causal meQTLs located more than 3 kb away from the methylated site.

We then explored the distribution of methylated sites that are associated with meQTLs in the context of other *cis*-regulatory annotations. Using the chromatin state annotations from Ernst et al. [50], we classified the genomic regions containing the assayed methylated sites as insulators, enhancers, or promoters (see Methods). Compared to the distribution of all assayed methylation sites, we found a relative depletion of sites associated with meQTLs at promoters (chi-square test; *P* < 10^−15^), and an enrichment of such sites at insulators (chi-square test; *P* < 10^−5^) and enhancers (chi-square test; *P* < 10^−9^; Table S1), consistent with previous work [8,40].

## QTLs for other regulatory phenotypes are often meQTLs as well

Our group has previously collected a number of genomic datasets from the same panel of Yoruba LCLs, pertaining to different regulatory mechanisms. We analyzed our methylation data in the context of these other data sets. We first performed a joint analysis of the methylation data with previously mapped eQTL data from the same LCLs [21]. We found that 146 (25%) of 595 eQTLs (classified at an FDR = 10%) within 3 kb of the genomic location of a methylation probe are also significantly associated with variation in DNA methylation (measured by the proximal probe; classified at an FDR = 10%). In other words, these SNPs are classified, using relatively stringent criteria, as both eQTLs and meQTLs (Fig. 2A). This represents a very strong enrichment of SNPs that are both eQTLs and meQTLs: the mean overlap expected by chance alone is 2.8% (*P* < 10^−5^; see Methods). Although we are unable to infer causality in this case (namely, to determine whether methylation patterns underlie gene expression levels or the other way around, or alternatively both phenotypes are responding to a third underlying factor), our observations indicate a substantial degree of coordination between methylation levels and gene expression.

**Figure 2.**
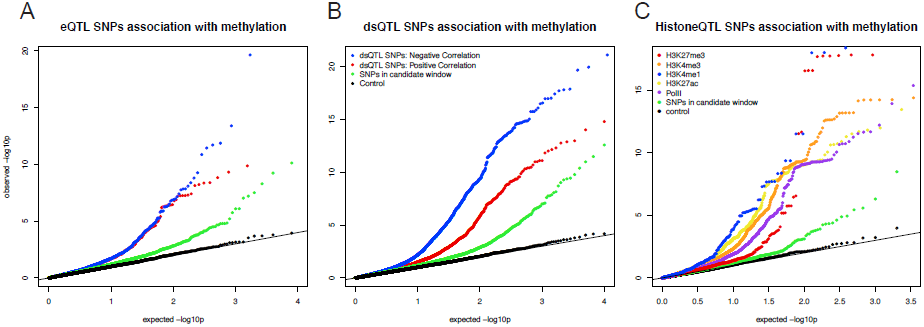
**A)** QQ plot of −log10 p-values for testing the null of no association between eQTL SNPs and methylation levels in sites within 3 kb. Positive correlations between expression and methylation levels are in red; Negative correlations are in blue, Data for random SNPs within the candidate window are in green; and data for a set of permuted genotype labels are in black. **B)** A plot of similar structure considering the associations of dsQTL SNPs [21] and with methylation levels at sites within 3 kb. **C)** A plot of similar structure considering the QQ plots of associations between histone modification QTLs [19] and methylation levels at sites within 3 kb.

Interestingly, roughly half of the sites classified as both eQTLs and meQTLs (70 of the 146 sites) are associated with positively correlated gene expression and methylation levels; namely, we observe a pattern whereby the genotypes that are associated with high expression levels are also quite often associated with high methylation levels. This pattern was observed both for methylation sites located within and outside gene bodies, yet we found that the CpG sites whose methylation levels are positively correlated with the expression levels of nearby genes are further from the gene’s TSS (median distance of 6,680 bp) than CpG sites whose methylation levels are negatively correlated with the expression levels of nearby genes (median distance of 1,020 bp; *P* = 0.018; Fig. S2). We were concerned that the more distal loci may be enriched for false positives. However, this observation remains significant (*P* = 0.027) even when we add effect size as a covariate in our model.

Next, we considered a joint analysis of the methylation data with QTL data for four histone modifications, PolII occupancy [19] and DNase I hypersensitivity profiles [21]. We found that QTLs associated with changes in any of these regulatory features are significantly more likely to also be associated with changes in methylation levels than expected by chance alone (by permutations; *P* < 10^−4^; Table 1; Fig. 2B, C). For example, 48% and 40% of QTLs associated with variation in H3K4me3 and H3K27ac, respectively, are also classified as meQTLs (at FDR = 10%). One particularly striking example of concerted changes in regulatory mechanisms that are associated with genetic variation at one locus is shown in Figure 3. The genotypes of a SNP located on chromosome 6, in an intron of the *HLA-DQB1* gene, are strongly associated with changes in DNase I hypersensitivity (*P* < 10^−9^), H3K4me3 (*P* < 10^−4^), H3k27ac (*P* < 10^−4^), gene expression levels (*P* < 10^−15^), and DNA methylation (*P* < 10^−10^).

**Table 1:**
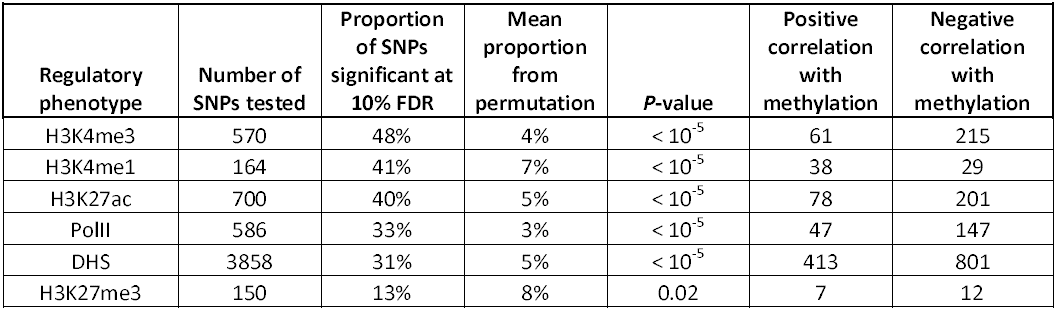
Associations between QTLs for other regulatory phenotypes and DNA methylation. For each regulatory phenotype we randomly sampled a matched number of SNPs, within 3 kb of a DNA methylation probe, 100,000 times. We calculated proportion of these tests significantly associated with methylation at an FDR of 10%. This was used to calculate the Max proportion from the subsample and the *P*-value columns.

**Figure 3.**
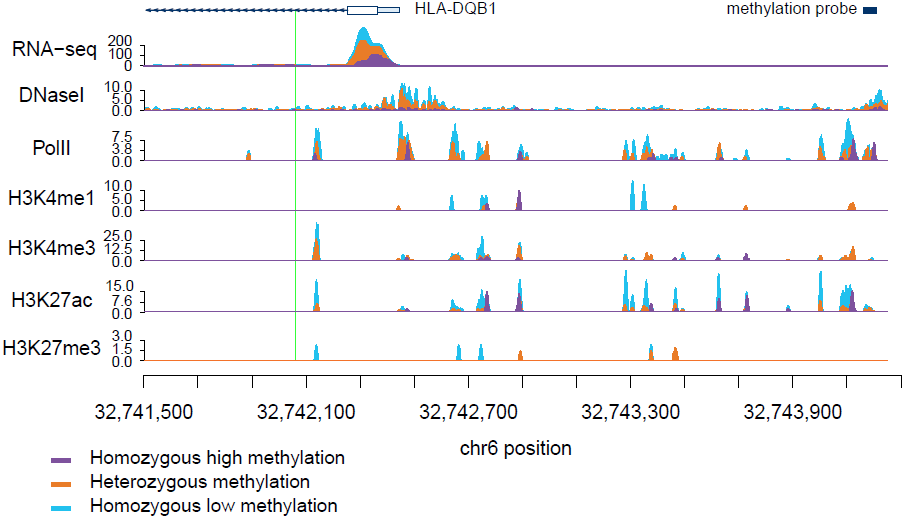
Read counts segregated by meQTL genotype for multiple regulatory phenotypes. The green line denotes the meQTL and the location of the probe measuring methylation data associated with the meQTL is identified by the black rectangle. The different colored data series indicate mean read depths segregated by genotype at the meQTL site: blue shows the homozygous genotype associated with low methylation level, orange shows the heterozygote, and purple the homozygous genotype associated with high methylation level. In this example, all of the regulatory phenotypes are negatively associated with DNA methylation levels.

Previous work has demonstrated that DNA methylation levels are generally negatively correlated with nearby levels of chromatin modifications associated with active transcription [12,41,51]. Yet, we found that methylation levels and chromatin features associated with active transcription are often positively correlated when variation in all features is associated in concert with a single QTL (Table 1; Fig. 2B, Fig. 3). It is important to note that often these regulatory regions, while proximal to each other, are not overlapping (eg. Fig. 4), suggesting a complex coordination across extended genomic regions.

**Figure 4.**
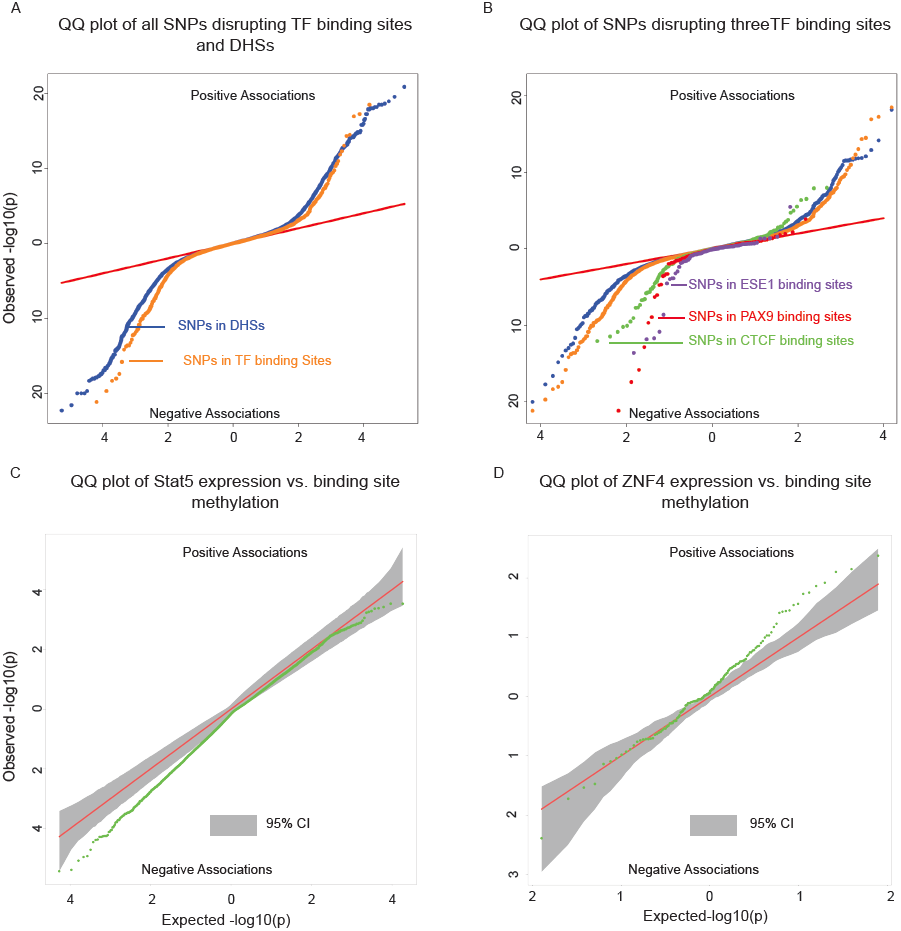
**A)** Two-sided QQ-plots describing the effect of TF binding on DNA methylation. For each SNP in a predicted TF binding site [52] we tested whether the SNP was associated with methylation at sites within 500bp. Positive associations (upper right quadrant) indicate that the allele associated with increased PWM score for the TF in question is associated with *increased* methylation; negative associations (lower left quadrant) indicate that increased PWM score is associated with *decreased* methylation. We used a random set of SNPs in DNase I hypersensitive sites (DHSs) to indicate the expected baseline. When considering the control DHS SNPs, the direction of the effects was chosen randomly for the purpose of plotting. Panel **B)** additionally highlights four TFs that show particular strong association with changes in methylation levels. **C)** Two-sided QQ-plot of associations between Stat5 expression and DNA methylation at sites within 500bp of Stat5 binding sites. **D)** QQ-plot of associations between ZNF274 expression and DNA methylation near ZNF274 binding sites. In both **C** and **D,** the grey shading indicates a region that would contain the data 95% of the time when the null hypothesis is true for all tests, obtained based on permutation of the expression data while holding the methylation data constant.

## Transcription factor binding may affect nearby patterns of DNA methylation

A major limitation of most genomic studies, including ours, is the difficulty of identifying casual mechanisms. However, we reasoned that we might be able to gain better insight about causality, or at least the likely order of events, if we focused on SNPs disrupting TF binding sites. It is reasonable to assume that the most direct outcome associated with such genetic variation is the disruption of TF binding. If these SNPs are also associated with changes in additional regulatory mechanisms, it might therefore be reasonable to further assume that changes in TF binding resulted in concerted changes in other regulatory phenotypes. Recent work has provided some measure of support for this rationale by suggesting that changes in TF binding can play causal roles in driving changes in histone marks [17–19] as well as DNase I hypersensitivity [21]. These results, in conjunction with previous examples of transcription factor binding altering methylation levels [34,44], led us to hypothesize that we could identify novel associations between TF binding and DNA methylation profiles. To do so, we examined the association of SNPs within TF binding with DNA methylation at nearby genomic regions.

To identify SNPs that are likely to directly affect TF binding we used DNase-seq data and the Centipede algorithm [52] to infer sites that are putatively bound by TFs in our LCLs. We next identified SNPs disrupting these putative binding sites and calculated a position weight matrix (PWM) score for each allele. We used SNPs that are in DNase I hypersensitive sites (DHSs) but not in known TF binding sites as a set of matched controls. Considering the data for all TFs together, we found that alleles with lower predicted TF binding affinity (i.e., lower PWM scores) are frequently associated with increased DNA methylation within 500bp of the binding site. The association was stronger than that observed for the control DHS SNPs (by permutations; *P* = 10^−5^; Fig. 4A). Considering binding sites for each TF separately, we identified three TFs (*CTCF*, *PAX9,* and *ESE1;* Fig. 4B), where a change in PWM score is significantly associated with the methylation level of probes within 500bp of the binding site (Table 2). Changes in the predicted binding efficiency of *ESE1* and *PAX9* are negatively associated with methylation levels, while changes in the predicted binding efficiency of *CTCF* are positively associated with methylation levels at some loci and negatively associated at others.

**Table 2:** Associations between SNPs disrupting TF binding sites and DNA methylation within 500pb of the binding site. For each TF we randomly sampled a matched number of SNPs, within any TF binding site with a CpGs less than 500bp away, 100,000 times. We calculated the proportion of tests significantly associated with methylation at an FDR of 10%. This was used to calculate the Max proportion from the subsample and the *P*-value columns.

| Name | SNPs tested | Proportion significant at 10% | Mean proportion from permutation | *P*-value |
| --- | --- | --- | --- | --- |
| CTCF | 370 | 15% | 3% | $< 10^{-5}$ |
| PAX9 | 85 | 11% | 3% | $< 10^{-4}$ |
| ESE1 | 55 | 15% | 2% | $< 10^{-4}$ |

Our observations indicate that the level of predicted TF binding is associated with variation in methylation levels near the binding site. Given this, changes in TF abundance (approximated by the estimated expression level of that TF) might also be associated with variation in methylation levels around the TF binding sites. To test this, we considered previously collected gene expression (RNA-seq) data from the same LCLs [21]. We found that the inter-individual variation in the expression levels of two TFs (*STAT5A* and *ZNF274*) is significantly correlated with variation in methylation levels around the TF binding sites (Fig. 4C/D). Specifically, an increase in *STAT5A* expression is associated with lower levels of DNA methylation and, interestingly, an increase in the expression of *ZNF274* is associated with increased levels of DNA methylation.

## meQTLs are enriched with loci associated with complex disease

Previous work has suggested links between DNA methylation, QTLs, and complex traits [31,53]. To further explore this in our data we used the NHGRI’s catalog of published genome-wide associations [54] to identify 391 SNPs associated with complex diseases that were within 3 kb of a methylation probe. We found that GWAS SNPs are significantly enriched among meQTLs (*P* < 10^−5^; Fig. S3); of the 2676 SNPs tested, 153 are also significantly associated with variation in methylation levels at an FDR of 10%. Given that LCLs are derived from B-lymphocytes and that DNA methylation exhibits tissue specificity, we hypothesized that the GWAS results would be enriched for genes pertaining to immune system functions. Using data from the original GWA studies we obtained a list of putatively affected genes associated with each of the 153 GWAS/meQTL SNPs. These genes are indeed enriched (FDR < 1.2%; Table 3) for KEGG pathways pertaining to immune function (eg. type 1 diabetes, antigen processing, autoimmune thyroid disease) and GO terms for immune function (eg. antigen processing and MHC class II receptor activity). We further found that genes implicated in the GWAS/meQTL analysis tend to be up regulated in peripheral blood leukocytes, compared to a background of multiple tissues (Table 3).

**Table 3:** DAVID analysis of meQTLs implicated in GWAS. The results from a DAVID analysis [66,67] of genes near a SNP whose genotype is associated with DNA methylation levels and complex disease. These data include GO terms, KEGG pathways, and up-regulated tissues.

| Category | Term | Fold Enrichment | Bonferroni | FDR |
| --- | --- | --- | --- | --- |
| KEGG_PATHWAY | Type I diabetes mellitus | 113 | 2.7E-11 | 3.9E-10 |
| KEGG_PATHWAY | Antigen processing and presentation | 101 | 1.9E-07 | 2.7E-06 |
| KEGG_PATHWAY | Graft-versus-host disease | 98 | 2.3E-07 | 3.3E-06 |
| KEGG_PATHWAY | Autoimmune thyroid disease | 91 | 3.4E-07 | 4.9E-06 |
| KEGG_PATHWAY | Allograft rejection | 88 | 4.1E-07 | 5.8E-06 |
| GOTERM_MF_FAT | MHC class II receptor activity | 190 | 2.0E-06 | 1.1E-05 |
| KEGG_PATHWAY | Cell adhesion molecules (CAMs) | 42 | 9.6E-07 | 1.4E-05 |
| KEGG_PATHWAY | Viral myocarditis | 69 | 1.4E-06 | 2.0E-05 |
| GOTERM_CC_FAT | MHC protein complex | 34 | 7.8E-06 | 5.9E-05 |
| UP_TISSUE | Blood | 10 | 1.0E-05 | 1.1E-04 |
| UP_TISSUE | Peripheral blood leukocyte | 47 | 1.6E-05 | 1.8E-04 |
| GOTERM_BP_FAT | Antigen processing and presentation | 25 | 3.8E-04 | 5.3E-04 |
| KEGG_PATHWAY | Asthma | 88 | 8.2E-04 | 0.012 |

## Discussion

Our study considered inter-individual variation in methylation profiles using LCLs. The LCL model is a somewhat artificial system, and indeed it has been previously demonstrated that the Epstein-Barr virus transformation of primary B cells into LCLs results in widespread DNA methylation changes [55,56]. However, it is also clear that a large number of B cell-specific characteristics remain in LCLs and, in general, important and insights regarding gene regulatory processes have been learned from studies in LCLs in particular, often by using a QTL mapping approach [55].

We have identified nearly 14 thousand CpG sites at which methylation levels are associated with genetic variation. The number and magnitude of associations are consistent with other recent meQTL studies of similar scale [40]. We took advantage of the fact that the LCLs we worked with are well studied (a clear advantage of the renewable LCL resource) to analyze the methylation data in combination with data on other regulatory mechanisms. We found strong evidence that DNA methylation is regulated in concert with other cellular phenotypes. Though the inference of causality is problematic for most genomic studies, including ours, we provided some indication that transcription factor binding may result in changes in DNA methylation patterns at nearby genomic regions.

Indeed, we found that, in general, SNPs disrupting TF binding sites are more likely to be associated with DNA methylation levels than SNPs within DNase I hypersensitive sites but not in TF binding sites. We believe that using SNPs disrupting putative TF binding sites provides a powerful way to re-examine the interplay between QTLs for regulatory phenotypes. Our observations therefore suggest that changes in the binding of *CTCF, PAX9, ESE1, STAT5,* and *ZNF274* result in changes in methylation patterns in nearby CpGs. This does not necessarily mean that the TF is directly regulating DNA methylation, but that changes in the binding of the TF (observed through change in mRNA abundance or PWM score) are the first step leading to a change in DNA methylation. In other words, our approach suggests that changes in TF binding are frequently a key early step in the regulatory cascade that leads to concerted changes in multiple mechanisms.

## The functional context of meQTLs

We observed an under-representation of meQTLs at promoters. We suggest two possible explanations for this observation; unfortunately, we currently lack the ability to distinguish between the two. First, a technical / statistical explanation: We may be underpowered to detect changes in methylation at promoters. We found that DNA methylation levels at promoters are, in general, less variable and have a lower average methylation level compared with other genomic regions, including enhancers (Fig. S4). The alternative explanation is more intriguing: It is possible that promoter methylation patterns are more often functional (with respect to their regulatory outcome) than methylation in other genomic regions. If so, promoter methylation patterns may evolve under stronger functional constraint, leading to lower true rates of meQTLs, as suggested previously [40].

Related to this interpretation, we have also shown that the relationship between DNA methylation and activating marks is more complex then previously appreciated. Negative correlations between DNA methylation levels and the expression of nearby genes have been observed frequently [8,12,33,35,57], but few have explored cases where DNA methylation is positively correlated with gene expression levels or activating chromatin marks [8,12,40]. When we examine joint QTLs, all regulatory phenotypes associated with active transcription exhibited an unexpectedly high proportion of positive correlations with methylation levels at nearby sites (Table 1). Previous work has shown that DNA methylation in gene bodies is often associated with activating histone modifications and increased expression levels [58,59], yet at least when we considered meQTLs, we did not observe a difference in the direction of correlations between CpGs within or outside gene bodies. Instead, we have found that when eQTL/meQTLs are positively correlated the respective TSS and CpG sites tend to be further from each other. These observations suggest that DNA methylation in more distal regulatory elements may be more likely to have an activating effect. This hypothesis is supported by the observed enrichment of CpG associated with meQTLs in enhancers and insulators, which are further from TSS than promoters.

We propose two alternative hypotheses to account for the observations of positive correlations between methylation and expression levels at nearly half of meQTLs/eQTLs sites. First, if the expression of a gene is tightly regulated, DNA methylation could serve as a fine-tuning tool. For example, over-activation by histone modifications could be suppressed using DNA methylation or vice versa. Indeed, while DNA methylation was considered a very stable epigenetic mark, recent work has demonstrated that DNA methylation levels can dynamically change *in vivo* on very fast (hours) time scales [60].

A second possibility is that observed positive correlations between methylation levels and the expression of nearby genes are due to 5-Hydroxymethylcytosine (5hMc), an additional modification to DNA methylation that has been implicated in the process of demethylation [61]. It has been shown that 5hMc has activating effects on transcription [62]. The bisulfite conversion approach we used does not allow us to distinguish 5hMc from DNA methylation. It is therefore possible that positive correlations between DNA methylation and expression or activating histone modifications are due to 5hMc.

## Summary

Our study joins a growing body of work, which indicates that methylation levels at a large number of loci across the genome are affected by genetic variation at nearby sites. In many cases, these meQTLs are also associated with variation in a variety of other types of chromatin changes, gene expression changes, and often - changes in disease risk. Our data is consistent with the notion that TF binding likely plays a role in altering methylation levels, but the mechanisms underlying the vast majority of meQTLs remain unclear. Similarly, we still do not understand in detail the mechanistic links between DNA methylation and other epigenetic marks and gene expression outputs, and these types of questions will no doubt be a fruitful area for future research.

## Materials and Methods

### DNA methylation array

To analyze DNA methylation, we extracted DNA from LCLs of 64 adult YRI HapMap individuals. The samples were bisulphite-converted and hybridized to the Infinium HumanMethylation450 BeadChip at the University of Chicago Functional Genomics facility. To validate the array probe specificity, probes were mapped to an *in silico* bisulfite-converted genome using the Bismark aligner [63]. Only uniquely mapped probes were retained (n = 459,221). We excluded probes on sex chromosomes (n = 14,016). Next, to eliminate the potential for spurious associations due to differences in probe hybridization affinity, we discarded probes (n = 104,720), overlapping known SNPs segregating in our panel based on our genotype data (see below). Following this series of exclusions, we kept data from 340,485 probes for subsequent analysis. Methylation levels are reported as β-values, which are considered estimates of the fraction of chromosomes methylated at a given site.

### Whole genome bisulfite sequencing

Bisulfite sequencing was performed using a modified version of the Illumina whole genome bisulfite sequencing protocol. Specifically, extracted DNA from LCL cell lines of 10 Yoruba HapMap population individuals and spiked-in unmethylated lambda phage DNA was fragmented into 100bp fragments using a Covaris ultra-sonicator. Fragmented DNA was blunt ended, repaired, and standard Illumin TruSeq adapters were ligated to the DNA fragments. DNA was then bisulfite-converted using the Invitrogen MethylCode Bisulfite Conversion Kit. The bisulfite-converted DNA was PCR amplified and sequenced using the Illumina HiSeq 2000. Each sample was sequenced in at least two lanes. Average genome-wide coverage ranged from 0.4x to 7.0x per sample with a median of 2.4x. Sequencing reads were trimmed for quality and to remove the adapter sequences. PCR duplicates were removed using the SAMtools software package. Reads were mapped using the Bismark aligner, which maps bisulfite converted DNA to a G to A and C to T converted human genome [63]. The bisulfite conversion efficiency was determined using the spiked-in lambda phage DNA. Conversion efficiency for all samples was estimated to be greater than 99%. Locus-specific methylation levels were estimated by obtaining the ratio of methylated to unmethylated CpG counts.

### Correlation of data from methylation array and bisulfite sequencing

To assess the overall agreement between the methylation array and the bisulfite-seq data we compared average methylation levels across CpG sites. To do so, we calculated the average of the untransformed array beta values from all 64 individuals at each CpG site, and compared these values to the estimated locus specific methylation level based on the sequencing data (by dividing the number of methylated reads by the total coverage of a given site in each individual, and calculating the mean across all individuals with at least 5 reads at that site). Correlation (Fig. S1) was assessed using the Spearman rank correlation (because the data are not normally distributed).

### Genotype data

We used the genotypes from a previous study of the same samples [21]. Briefly, genotypes were obtained by combining and imputing genotype based on the 1000 Genomes Project and HapMap [48,49]. A reference panel was built using all 210 YRI individuals (excluding 1^st^ degree relatives). If genotypes were available from multiple datasets the dataset that was expected to be most accurate on average was chosen (1000 Genomes high coverage, followed by HapMap, then 1000 Genomes low coverage, respectively). This reference panel was used to impute missing genotypes for individuals in our cohort using the BIMBAM software [64]. Genotype information was obtained for roughly 15.8 million variants genome-wide. The genotypes that we used can be found at http://eqtl.uchicago.edu/Home.html.

### QTL analysis

The distribution of methylation array data is non-Gaussian. We therefore quantile-normalized the data to a standard normal first, across all probes within an individual, and then across all individuals at each probe. We tested for confounders using principal component analysis. No known confounders were significantly correlated with a PC (Fig. S7). However, we found that removing four PCs provided optimal power to detect meQTLs (Table S2). We then identified meQTLs by testing (using standard linear regression) for associations between normalized methylation levels and genotypes at all SNPs that were within 3 kb of an assayed CpG. We only tested SNPs with a minor allele frequency greater than 5%. An FDR was computed using the R-package qvalue [65]. To investigate the overlap between QTLs for other molecular phenotypes and meQTLs we identified SNPs previously associated with changes in histone modifications, PolII, DHS, expression and complex diseases (using GWAS results) [19,21,54]. The rationale for this analysis is that the observation that a SNP is a QTL for other traits increases the overall likelihood that the SNP may also be associated with changes in methylation levels (in other words, we use previous observations as priors). Significant QTLs for any of the tested regulatory phenotypes or complex diseases, that were located within 3 kb of a methylation probe, were then tested for association with methylation levels. For each class of previously identified QTLs an independent FDR [65] was calculated to assess the significance of association with methylation levels.

To ensure that our results are not markedly impacted by the choice of normalization procedure, we also considered two alternative approaches. First, the data were quantile-normalized to a standard normal across all probes within an individual. This approach resulted in a minor excess of small p-values in the QTL analysis of permuted data (Fig. S5). Second, we quantile-normalized data from a given probe to a standard normal across all individuals. This method resulted in considerable variation in mean methylation levels across individuals, which is not ideal since the variable means may reflect array variation rather than true biology. Regardless of the specific properties (and possible shortcomings) of the alternative normalization and data processing approaches, the majority of meQTL associations we report remained significant (8,684 without removing PCs, 8863 when normalized by individual, 5496 when normalized by probe, and 6283 when the data were untransformed; Fig. S6).

### Aggregation of bisulfite sequencing data

We used the bisulfite sequencing data to generally validate the meQTLs identified using the array data, and more importantly, to visualize the association of meQTLs with methylation levels at CpGs that are located near each other. Since the sequence data are sparse (because the coverage is low) and available for only a small number of individuals, we only considered an aggregate analysis across all individuals and across all the previously identified meQTL associated CpGs. Specifically, for each meQTL we separated the sequenced individuals by genotype (i.e., the genotypes associated with high methylation levels, heterozygote, or those associated with low methylation levels). Next, we counted the number of methylated and unmethylated reads in 51bp windows sliding across a 5 kb region centered on the associated CpG for each meQTL. The mean aggregate methylation levels for each window position and each genotype class were calculated as the sum of the number of methylated reads divided by the sum of total reads for that window and genotype class. We averaged this estimate across all meQTLs genome-wide. The result is an aggregate plot of the average methylation levels by genotype class, showing the spatial distribution of CpG methylation in a 5 kb window (Figure 1B).

### Identification of candidate causal SNPs from meQTL data

Due to LD, the causal site for any given meQTL is typically ambiguous. In addition, though we used 1000 genome sequence data and imputation, we expect that a subset of common SNPs are missing from our data. For this reason, it is challenging to obtain an accurate estimate of the distribution of distances between probes and causal meQTL sites. In previous work, our group tackled this problem using a Bayesian model [12]. Here, since we have a much larger number of meQTLs (then eQTLs or dsQTLs, for example), we focused on a set of meQTLs where there is a single clear candidate variant that is likely to drive the signal. Specifically, we identified meQTLs for which the p-value of the most significant SNP is at least two orders of magnitude lower than that of the next most significant SNP (within a slightly larger, 10 kb window). Previously, we used simulations to show that these stringent criteria provide strong enrichment for causal sites [7]. In reality, we consider these sites as putatively causal because the evidence supporting their role is circumstantial.

### Inclusions of previous data collected from the same samples

DNase-seq data for 70 individuals, ChIP-seq data for 10 individuals and RNA-seq data for 69 individuals were obtained from previous studies performed in our labs [2,19,21]. In Figure 3, mapped fragments are reported as fragments per kilobase per million mapped reads (FPKM) and are smoothed using a 21bp Savitzky-Golay filter.

### Association between transcription factor binding and DNA methylation

We performed analysis that focused on SNPs that disrupt TF biding sites. To do so, we used inferences of TF binding based on DNase I sequencing data that were obtained from a previous study [21], which applied the Centipede algorithm [52] to DNase-seq data from the same LCLs. We identified putative binding sites overlapping genetic variants and calculated a position weight matrix (PWM) score for both alleles at each locus. Linear regression was then performed to identify associations between the PWM scores of each genotype and the methylation levels of CpGs within 500 base pairs of the motif position.

### Association between transcription factor expression levels and DNA methylation at CpGs near the TF binding sites

RNA-seq data for 56 of the 64 individuals with methylation array data were obtained from Degner et al. [21]. The mRNA levels of the transcription factors were standardized to RPKM and then quantile normalized. We used ChIP-seq broad-peak calls for 100 TFs, measured by the ENCODE project in the lymphoblastoid cell line GM12878, to identify TF binding sites [14]. (These data were downloaded from the ENCODE website (http://encodeproiect.0rg/ENCODE/) in July 2013). If the TF ChIP-seq was performed in multiple replicates, only the peaks found in all replicates were considered as binding sites. A Pearson correlation test was performed between the TF expression and DNA methylation levels measured by probes within 500 base pairs of TF binding sites. Given our expectation that TF expression would have a *trans* effect on DNA methylation genome-wide, we anticipated removing PCs from the methylation data would diminish our ability to identify associations. Indeed we find that using data with PCs removed reduces our power to identify associations. As such, we used methylation data that had only been normalized (first by individual then by probe) for this analysis.

### Pathway analysis of GWAS associated genes

We performed a pathway analysis of GWAS associated genes using the DAVID program [66,67]. DAVID allows the user to input a custom “background” set of genes from which the program computes a null hypothesis. Since there is a known bias toward immune system genes in GWA studies we used all genes implicated in GWA studies as our “background”. Thus, observed significant enrichments are beyond the bias in GWAS results.

## Acknowledgements

We thank the three anonymous reviewers for helpful comments on our manuscript, members of the Pritchard, Stephens and Gilad laboratories for many valuable discussions, and the ENCODE Project for publicly available ChIP-seq data.

## Accession Numbers

Data from the methylation array and bisulfite sequencing are available at the GEO database (accession number GSE57483). A summary table of the meQTLs is available at the Gilad lab website http://giladlab.uchicago.edu/Data.html.

## Figure Legends

**Figure S1.** Scatterplot of CpG methylation levels estimated from the Illumina array and from whole genome bisulfite sequencing.

**Figure S2.** A boxplot of distances from methylation probe to transcription start site for eQTL/meQTLs. The boxplot on the left represents QTLs where methylation and expression are negatively correlated. The boxplot on the right represents QTLs where methylation and expression are positively correlated.

**Figure S3.** QQ-plot of associations between SNPs implicated in GWAS studies and DNA methylation. The red points are all SNPs from GWAS studies within 3 kb of a methylation probe. The black points are a subsample of all the SNPs within 3 kb of a methylation probe.

**Figure S4.** Distributions of methylation levels at array probes in promoters and enhancers, respectively, and the full distribution across all probes. Promoters have reduced variability compared to all probes and to enhancers.

**Figure S5.** QQ-plot of all SNPs within 3 kb of a methylation probe normalized by either **A)** individual or **B)** probe. Visible inflation of associations is observed when normalizing by individual.

**Figure S6.** The T-statistics of meQTLs identified in this study when regression is performed using other array normalization strategies. The histograms show the absolute T-statistic for **A)** untransformed data, **B)** data normalized by individual only, **C)** data normalized by probe only, and **D)** normalized by individual then probe. The blue histogram represents permuted genotypes (controls) and the red histogram represents the meQTLs.

**Figure S7**. PCA plots showing the first two PCs separated by **A)** sex, **B)** bisulfite conversion batch, or **C)** array batch. None of the known potential confounders are associated with PC1 or PC2. PC1 explains roughly 8% of the variance.

**Table S1.** The data used to test for enrichments/depletions of probes measuring methylation levels meQTL associated CpGs. The first column is the number of meQTLs associated CpGs within the specified genomic feature (eg. promoter). The second column is the total number of probes within the specified genomic feature. To calculate the chi-square statistic a two by two contingency table was created using the first two columns (described above), the total number of meQTL associated CpGs, and the total number of probes on the array.

**Table S2.** This table shows the differing number of meQTLs identified (at an FDR of 10%) after removing varying numbers of PCs.

